# TTBK2 and primary cilia are essential for the connectivity and survival of cerebellar Purkinje neurons

**DOI:** 10.1101/689448

**Authors:** Emily Bowie, Sarah C. Goetz

## Abstract

Primary cilia are vital signaling organelles that extend from most types of cells, including neurons and glia. However, their function, particularly on neurons in the adult brain, remains largely unknown. Tau tubulin kinase 2 (TTBK2) is a critical regulator of ciliogenesis, and is also mutated a hereditary neurodegenerative disorder, spinocerebellar ataxia type 11 (SCA11). Here, we show that conditional knockout of *Ttbk2* in adult mice results in degenerative cerebellar phenotypes that recapitulate aspects of human SCA11 including motor coordination deficits, loss of synaptic connections to Purkinje cells (PCs), and eventual loss of PCs. We also find that the *Ttbk2* conditional mutant mice quickly lose cilia throughout the brain. We show that conditional knockout of the key ciliary trafficking gene *Ift88* in adult mice results in nearly identical cerebellar phenotypes to those of the *Ttbk2* knockout, supporting disruption of ciliary signaling as a key driver of these phenotypes. Our data suggest that primary cilia play an integral role in maintaining adult neuronal function, and offers novel insights into the mechanisms involved in neurodegeneration.

## INTRODUCTION

Primary cilia function as compartments that mediate and integrate essential signaling pathways. Because of their critical roles in developmental signaling pathways including Hedgehog (HH) signaling (Goetz and Anderson, 2010), disruptions to cilium assembly, structure, or function are associated with a number of hereditary developmental syndromes, which are collectively termed ciliopathies. Among the more common pathologies associated with ciliopathies are a variety of neurological deficits (Reiter and Leroux, 2017). During development, ciliary signals drive the proliferation and patterning of neural progenitor populations (Guemez-Gamboa et al., 2014). Cilia then persist on post-mitotic neurons during development and through adulthood (Sterpka and Chen, 2018). However, we have only an incomplete understanding of the roles for primary cilia on differentiated, post-mitotic neurons, particularly within the adult brain.

Nevertheless, newly emerging evidence suggests that cilia and ciliary signaling may be important in this context: Cilia are required for the establishment of synaptic connectivity in hippocampal dentate granule neurons (Kumamoto et al., 2012) and in striatal interneurons (Guo et al., 2017). Neuronal cilia also concentrate a wide array of GPCRs and other neuropeptide and neurotrophin receptors important for complex neurological functions (Berbari et al., 2008; Domire et al., 2011; Green et al., 2012; Guadiana et al., 2016). Defects in cilia structure have also been associated with number of neurodegenerative and neuropsychiatric conditions(Chakravarthy et al., 2012; Dhekne et al., 2018; Keryer et al., 2011; Muñoz-Estrada et al., 2017). Huntington’s disease (HD) is caused by CAG repeat expansions in the gene *huntingtin*, which encodes a microtubule-associated protein (HTT) that localizes to the centrosome(Saudou and Humbert, 2016). Pathogenic HTT protein in transgenic mouse models is associated with abnormally long primary cilia (Keryer et al., 2011). Additionally, the Parkinson’s disease-associated kinase, LRRK2, is linked to ciliary dysfunction. Human disease-associated mutations to LRRK2 interfere with cilia formation in cultured cells and in mouse models through regulation of RAB8 and RAB10 by LRRK2 (Dhekne et al., 2018).

More directly, we have shown that Tau tubulin kinase 2 (TTBK2), a kinase causally mutated in the hereditary neurodegenerative disorder SCA11 (Houlden et al., 2007), is an essential regulator of ciliogenesis (Goetz et al., 2012). These mutations are frameshift-causing indels that result in premature truncation of TTBK2 at ∼AA 450. SCA11 is characterized by the loss of Purkinje cells (PC) in the cerebellum, causing ataxia and other motor coordination deficits(Houlden et al., 2007; Seidel et al., 2012). Recently, we demonstrated that SCA11-associated alleles of *Ttbk2* act as dominant negatives, causing defects in cilium assembly, stability, and function (Bowie et al., 2018).

Given the association between the SCA11-associated truncations to TTBK2 and ciliary dysfunction, we have set out to test whether loss of TTBK2 function within the adult brain is associated with degeneration of cerebellar neurons. The cerebellum is the region of the brain responsible for controlling motor coordination, learning, and other cognitive functions (Purves et al., 2001). The development and morphogenesis of the cerebellum depends on primary cilia, which are critical for the expansion of granule neuron progenitors (Chizhikov et al., 2007; Spassky et al., 2008). PCs, granule neurons and interneurons as well as Bergmann glia (BG) are ciliated in the adult cerebellum as well as during development. However, the roles of cilia and ciliary signaling in the adult cerebellum are unknown.

In the present study, we show that conditional loss of *Ttbk2* during adulthood as well as genetic targeting of cilia using an *Ift88* conditional mutant cause similar degenerative changes to the cerebellar architecture that worsen over time. These cellular changes are accompanied by motor coordination phenotypes in the mice. We demonstrate that loss of *Ttbk2* and cilia leads to altered Ca^++^ homeostasis in PCs, and ultimately to loss of these neurons. We provide strong evidence that primary cilia and ciliary signals are important for maintaining the connectivity of neurons within the brain, and we provide the first evidence that dysfunction of primary cilia can cause or contribute to neurodegeneration within the mammalian brain.

## RESULTS

### Loss of *Ttbk2* from the adult brain causes SCA-like cerebellar phenotypes

Mutations within *Ttbk2* cause the adult-onset, neurodegenerative disease SCA11; however the etiology of SCA11 is ill-defined. SCA11 is somewhat unusual among spinocerebellar ataxias in part because the reported causal mutations are base pair insertions or deletions(Houlden et al., 2007; Johnson et al., 2008; Lindquist et al., 2017), rather than CAG repeats which is the genetic cause of most SCAs(Hersheson et al., 2012). While mice heterozygous for a knockin of one of the human-disease associated SCA11 mutations do not exhibit degenerative phenotypes(Bouskila et al., 2011; Bowie et al., 2018), we postulated based on our previous studies that a knockout of *Ttbk2* from the adult mouse brain might cause neurodegenerative phenotypes that recapitulate SCA11. To test this, we obtained a conditional allele of *Ttbk2* (*Ttbk2^tm1c(EUCOMM)Hmgu^*) from the European Mutant Mouse Cell Repository, (referred to from here as *Ttbk2^fl^*). We then crossed *Ttbk2^fl^* mice to a mouse line expressing tamoxifen-inducible Cre recombinase driven by a ubiquitously expressed promoter, *UBC-Cre-ERT2(Ruzankina et al., 2007)*. Using this model, we induce recombination of *Ttbk2* in all tissues of the mouse, including the brain, upon injection with tamoxifen (TMX). Because development of the mouse cerebellum is complete by P21(Marzban et al., 2014), we chose this time to begin our TMX injections. For all of our experiments, control animals are either siblings with the same genotype (*Ttbk2^fl/fl^;UBC-Cre-ERT2*^+^) injected with oil vehicle only, or *Ttbk2^fl/fl^;UBC-Cre-ERT2^-^* sibling mice injected with the same dose of TMX.

To assess the consequences of loss of *Ttbk2* on the adult brain, we examined *Ttbk2^fl/fl^;UBC-Cre-ERT2^+^,*TMX treated animals (referred to from here as *Ttbk2^c.mut^*) at 4 months of age (3 months post TMX). The brains of *Ttbk2^c.mut^* mice have slightly smaller olfactory bulbs, but the overall gross morphology of the cortex and cerebellum was unchanged (Figure S1A). Since human mutations in *Ttbk2* cause SCA11(Houlden et al., 2007; Johnson et al., 2008; Lindquist et al., 2017), we examined the architecture and connectivity of neurons within the cerebellum to assess whether the *Ttbk2^c.mut^* animals exhibited phenotypes similar to those described for mouse models of other subtypes of SCA. Within the cerebellum, PCs are the major source of functional neuronal output(Cerminara et al., 2015). PCs receive excitatory inputs primarily from parallel fibers and climbing fibers. Parallel fibers extend from the granule neurons, which are the population of densely packed neurons found directly beneath PCs(Marzban et al., 2014). Climbing fibers extend from neurons of the Inferior Olivary Nuclei (ION) in the medulla of the brain stem(Purves et al., 2001). These connections are essential for PC functional output, and the dysfunction or loss of these connections, particularly the VGLUT2+ synapses from the climbing fibers, has been shown in various mouse models of SCA to be linked to pathology and disease progression(Duvick et al., 2010; Ebner et al., 2013; Furrer et al., 2013; Smeets and Verbeek, 2016; Smeets et al., 2015).

Within 3 weeks following induction of recombination with TMX, *Ttbk2^c.mut^* mice exhibited apparent locomotor deficiencies when observed in their cage (Supplemental Video S1). To further examine motor coordination in *Ttbk2^cmut^* mice, we employed a rotarod performance test. *Ttbk2^c.mut^* mice exhibited a shorter latency to fall compared to the littermate controls in each trial, for both the accelerating rotarod analysis as well as the steady speed rotarod analysis (Figure 1A, 1B). These results indicate that *Ttbk2^c.mut^* mice are impaired in their motor coordination after the loss of *Ttbk2* throughout the adult mouse brain, consistent with motor deficits by observed in multiple mouse models of SCA(Lalonde and Strazielle, 2019)(Klockgether et al., 2019).

**Figure 1.**
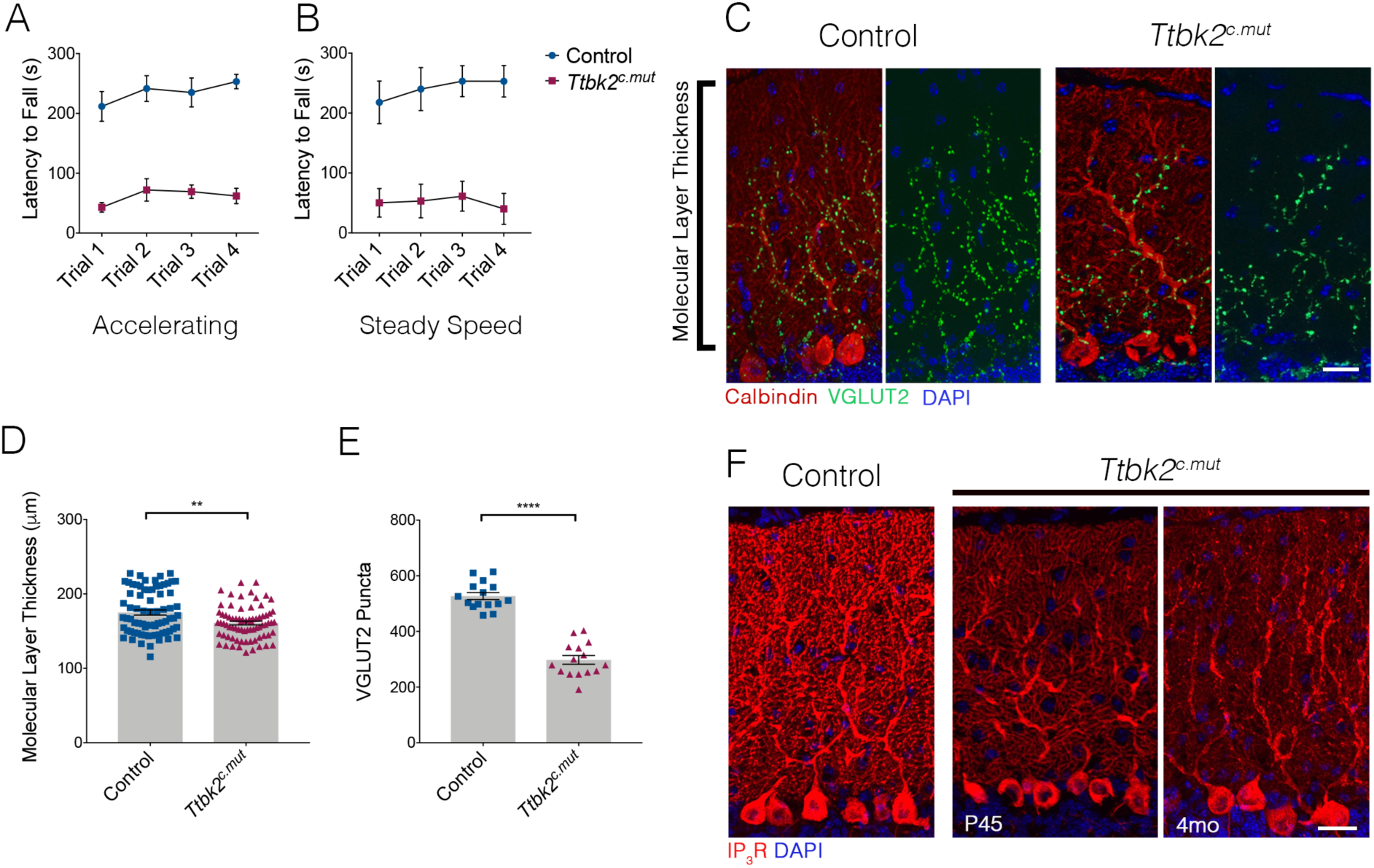
Loss of *Ttbk2* causes SCA-like phenotypes. **(A, B)** Accelerating and Steady Speed Rotarod Performance Test between *Ttbk2^c.mut^* and littermate controls. *Ttbk2^c.mut^* animals have a shorter latency to fall time in both tests, indicative of impaired motor ability (n=9 biological replicates for Control, 8 biological replicates for *Ttbk2^c.mut^*). **(C)** Cerebellar tissue from control and *Ttbk2^c.mut^* mice at three months after loss of *Ttbk2*, immunostained for Calbindin to label Purkinje cells (red) and VGLUT2 to show climbing fiber synapses (green). *Ttbk2^c.mut^* animals show a reduction in VGLUT2 positive synapses throughout the cerebellum three months after loss of TTBK2. Scale bar = 50μm **(D)** Quantification of molecular layer length in *Ttbk2^c.mut^* cerebellar tissue (each point represents one measurement, 75 measurements overall. n=3 biological replicates. p=0.0011 by student’s unpaired t-test, error bars indicate SEM 14.12 +/- 4.254) **(E)** Quantification of VGLUT2+ puncta throughout PC dendrites. *Ttbk2^c.mu^*^t^ animals show a significant reduction in these VGLUT2+ synapse terminals (each point represents one measurement, 15 measurements per genotype, n=3 biological replicates. p<0.0001 by student’s unpaired t-test, error bars indicate SEM 229.5 +/- 20.15). **(F)** Immunostaining for IP3R to label calcium channel abundance (red) and nuclei (blue). Loss of IP3R expression is seen as early as P45 in *Ttbk2^c.mut^* cerebellum. By four months, IP3R expression is no longer localized to secondary dendrites throughout the dendritic tree of PCs in four month old *Ttbk2^c.mut^* animals. Scale bar = 50μm.

At 4 months of age, the number of PCs is not affected in *Ttbk2^c.mut^* animals. However, we observed thinning of the molecular layer of the cerebellum, which is comprised of the elaborate dendrites extended from the PCs (Figure 1C, 1D. 175μm +/- 3.422 for Control vs. 160.9μm +/- 2.527 for *Ttbk2^c.mut^*). More dramatically, upon examination of the synaptic marker VGLUT2, which marks presynaptic terminals between PC dendrites and climbing fibers from the ION, we found a marked reduction in these puncta throughout the *Ttbk2^c.mut^* cerebellum compared to controls (Figure 1A, 1E 527.3 puncta +/- 12.68 for Control vs. 297.8 puncta +/- 15.65 for *Ttbk2^c.mut^*). Loss of TTBK2 protein was confirmed with western blot analysis on cerebellum lysates from *Ttbk2^c.mut^* animals and littermate controls (Figure S1B). We found no phenotypic differences between control condition animals or pre-induction *Ttbk2^fl/fl^;UBC-Cre-ERT2+* animals (Figure S1C, S1D, S1E). This loss of climbing fiber synapses we observed when TTBK2 is removed in adulthood is reminiscent of phenotypes observed in SCA1 as well as SCA7(Duvick et al., 2010; Furrer et al., 2013), indicating that *Ttbk2^c.mut^* mice exhibit degenerative cerebellar phenotypes recapitulating other subtypes of SCA.

### Calcium modulation is impaired after cilia loss

In order for PCs to function, they use an intracellular calcium modulation network. Inositol 1,4,5-trisphosphate receptors (IP3Rs) are key calcium channel regulators within this network, needed for calcium release from the surrounding ER throughout the PC (Sarkisov and Wang, 2008). Precise regulation of IP3R activity is critical, and a delicate balance of calcium channel release is imperative to the overall function of the cerebellum. Mutations in the IP3R1 gene have been linked to SCA15 and SCA29, while overexpression of IP3R1 is causes phenotypes within SCA2 and SCA3 (Tada et al., 2016). Due to these links to other SCA-related phenotypes, we examined IP3R expression throughout the *Ttbk2^c.mut^* animals. We found that IP3R expression is reduced in *Ttbk2^c.mut^* compared to controls starting at P45, with expression being strongly reduced in the dendrites by four months of age (Figure 1F). Taken together, this data suggests that the motor coordination phenotypes revealed by the rotarod performance test can in part be explained by the PCs inability to properly modulate calcium levels throughout the cerebellum due to loss of this receptor at the PC dendrites.

### Loss of *Ttbk2* causes changes to ION neurons and BG

While our data indicates that *Ttbk2* is essential to maintain the connectivity of PCs, we also tested whether other key cell types within the hindbrain that have been linked to the pathology of SCA are affected. Climbing fibers extend from neurons of the ION in the medulla. These fibers traverse the brain stem, enter the cerebellar cortex, and innervate the PC dendrites (Watanabe and Kano, 2011). Since we saw a reduction of the synapses between these climbing fibers and PC dendrites indicated by a reduction in VGLUT2 synapses, we examined the soma of the ION neurons from which these climbing fibers extend from. In several subtypes of SCA, including SCA1, 2, 3, 6 and 7 (Seidel et al., 2012), the pathology of the disorder is characterized in part by the shrinking of the ION soma; a characteristic that is also observed in mouse models of these diseases. Neurons within the ION can be identified by the dual expression of Calbindin and NeuN in the medial ventral region of the medulla (Figure S2A, S2B). We used NeuN to label the perikaryon of neurons within the ION, and measured the area of the somata of ION neurons, comparing *Ttbk2^c.mut^* and control animals. We found a significant reduction in the area of ION neuron soma in *Ttbk2^c.mut^* mice at 4 months of age compared to controls (Figure 2A, 2B; 180.4μm^2^ +/- 1.93 for Control vs. 109μm^2^ +/- 1.01 for *Ttbk2^c.mut^*).This implies that, in addition to the PCs themselves, the neurons sending critical inputs to the PCs are perturbed in *Ttbk2^cmut^* mice.

**Figure 2.**
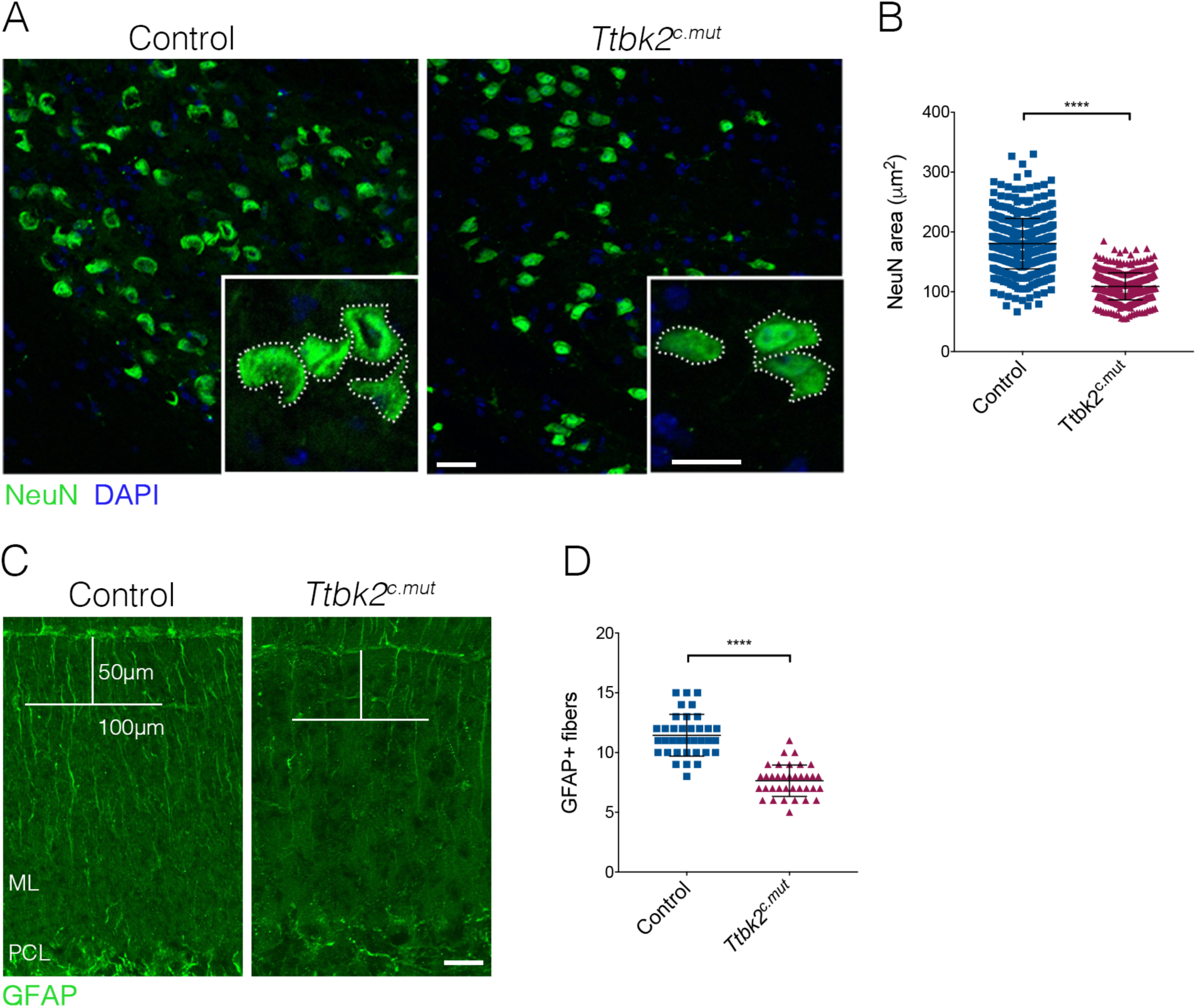
Loss of *Ttbk2* results in changes to a variety of cerebellar cells. **(A)** Representative images of neurons in the inferior olivary nucleus (ION) located in the medulla. Neurons are immunostained with NeuN (green). Insets show how area was measured. Scale bar = 50μm (20μm inset) **(B)** Quantification of NeuN area. ION neurons have reduced area in *Ttbk2^c.mut^* animals compared to control (each point represents single cell measurement of which >150 measurements were made per replicate. n= 3 biological replicates, p<0.0001 by unpaired student’s t-test, error bars indicate SEM 71.43 +/- 2.16). **(C)** GFAP staining showing Bergmann glial fibers throughout the molecular layer. In *Ttbk2^c.mut^* animals, density of these fibers is reduced. Quantification was done as previously described (Furrer et al., 2011) in which a 50μm line was drawn from the pial surface of the folia, and a 100μm across. Glial fibers that fully crossed the 100μm line were scored. Scale bar = 20μm. **(D)** Quantification of GFAP+ glial fibers which crossed the 100μm line (each point represents an image quantified, 36 images quantified per genotype across n=3 biological replicates. p<0.0001 by unpaired student’s t-test, error bars indicate SEM 3.80 +/- 0.36).

Throughout the brain, astrocytes and glia also play important roles in maintaining synaptic connectivity and strength. In the cerebellum, the processes of the BG are interspersed with PC dendrites in the molecular layer, with BGs enwrapping the excitatory synapses of the PCs (Leung and Li, 2018). Since defects in BG morphology have been linked to the etiology of SCA7, we investigated the BG population in *Ttbk2^c.mut^*. To assess the morphology of BGs in the *Ttbk2^cmut^* animals and evaluate whether defects in these cells may contribute to the phenotype. We used GFAP to visualize BG fibers which extend throughout the cerebellar folia. We found that the numbers of glial fibers were moderately reduced in *Ttbk2^c.mut^* cerebellar folia compared to littermate controls(Fig. 2C, 2D; 11.44 BG fibers +/- 0.29 for Control vs. 7.64 BG fibers +/- 0.22 for *Ttbk2^c.mut^*), suggesting that loss of *Ttbk2* affects the morphology, and perhaps the function of these cells. Taken together, this data suggests that loss of *Ttbk2* affects several cell types throughout the cerebellum as well as within the medulla, underscoring the widespread importance of *Ttbk2* within these tissues.

### Non-cell-autonomous requirements for *Ttbk2* in PCs to maintain their connectivity

Dysfunction and eventual atrophy of the PCs in the cerebellum is the primary pathology underlying SCA11 in human patients (Houlden et al., 2007). In our conditional *Ttbk2* mutant mice, the most prominent phenotype is altered connectivity of the PCs with additional cellular changes seen in the BGs as well as ION neurons. To determine the degree to which these defects are due to non-cell-autonomous requirements for *Ttbk2* in the PCs versus other cell types that make up the cerebellar circuit, we used the PC-specific Cre line *PCP2-Cre*. *PCP2-Cre* drives recombination specifically in PCs within the cerebellum beginning at P6 (Zhang et al., 2004). At P30, *Ttbk2^fl/fl^;PCP2-Cre^+^ (*referred to from here as *Ttbk2^PCP2^)* animals have normal cerebellar structure and synapse connectivity, with molecular layer thickness that is comparable to littermate control animals (Figure 3A, 3C. 202.7μm +/- 3.51 in P30 control vs. 191.5μm +/- 3.21 in P30 *Ttbk2^PCP2^*). Synapse connectivity in the PC by measure of VGLUT2 synapses is not significantly changed between P30 control and *Ttbk2^PCP2^* animals (Figure 3A, 3D. 548.2 puncta +/- 13.36 in P30 control vs. 538.9 puncta +/- 18.14 in P30 *Ttbk2^PCP2^*). This indicates that despite loss of *Ttbk2* during PC postnatal development, initial connections between PCs and climbing fibers are established normally. By P90, *Ttbk2^PCP2^* animals exhibited phenotypes largely recapitulating those observed in the *Ttbk2^cmut^* animals. While molecular layer thickness between the P90 Control and *Ttbk2^PCP2^* was not changed (Figure 3B, 3C. 202.9μm +/- 2.11 in P90 control vs. 197.4μm +/- 4.18 in P90 *Ttbk2^PCP2^*), we see a significant decrease in VGLUT2 puncta throughout the cerebellum (Figure 3B, 3D. 549.9 puncta +/- 15.47 in P90 control vs. 476.9 puncta +/- 15.82 in P90 *Ttbk2^PCP2^*), indicating that PCs have started to lose these important connections from the climbing fiber synapses.

**Figure 3.**
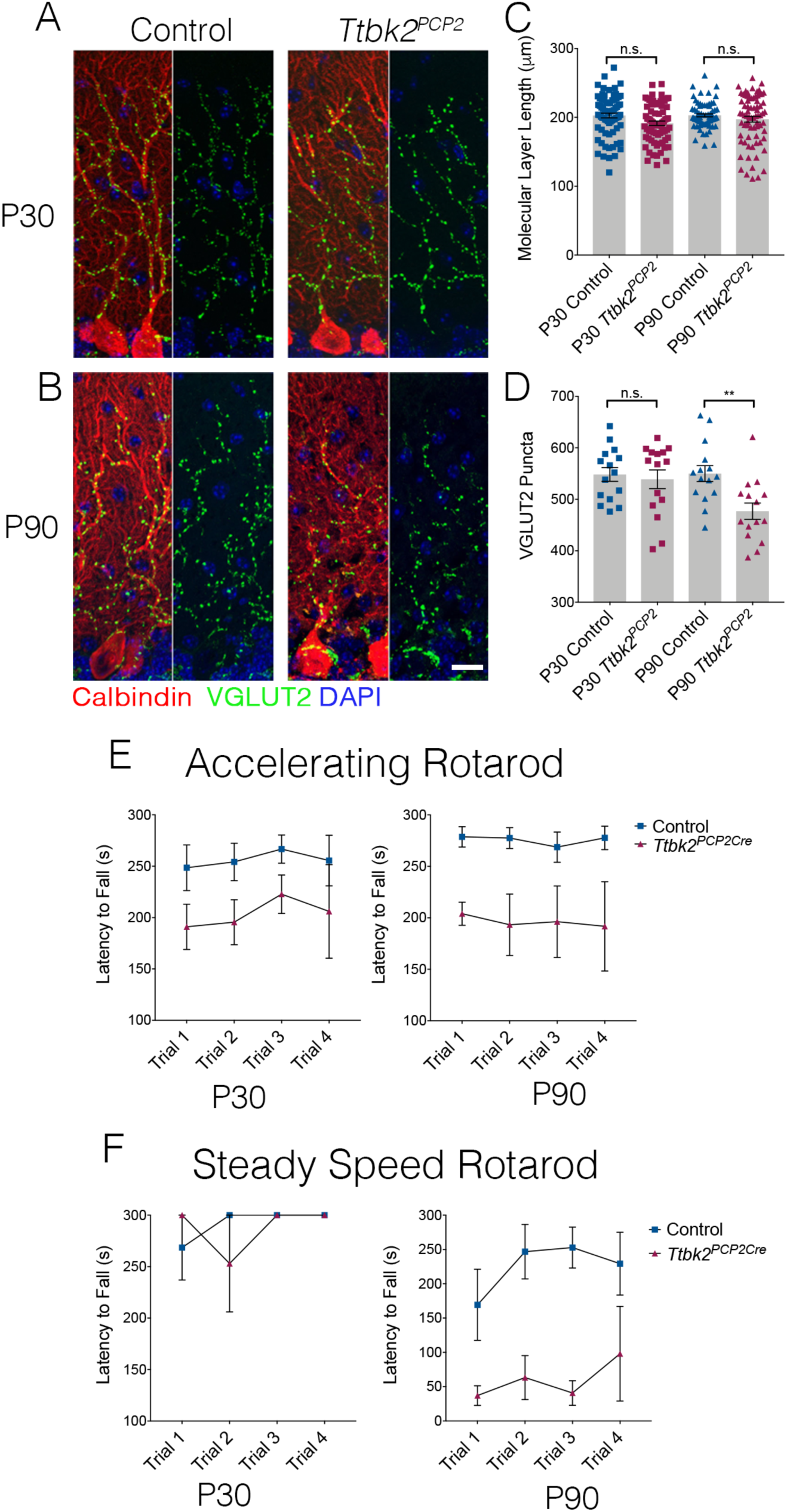
Cell autonomous requirements for *Ttbk2* in PCs. **(A,B)** Representative images of Control and Ttbk2^f/f^;PCP2Cre+ (*Ttbk2^PCP2^*) animals at age P30 (**A**) and P90 (**B**), immunostained for Calbindin to label PCs (red), VGLUT2 to label synapses (green), and nuclei (blue). VGLUT2 terminals are reduced in P90 *Ttbk2^PCP2^* animals compared to P30 *Ttbk2^PCP2^* animals. Scale bar = 20μm. (C) Quantification of molecular layer length in P30 and P90 *Ttbk2^PCP2^* and control animals (each point represents one measurement, 75 measurements per genotype. n=3 biological replicates. No significance reported by One-way ANOVA with Tukey correction, error bars indicate SEM. **(D)** Quantification of VGLUT2+ puncta analysis in P30 and P90 *Ttbk2^PCP2^* and control animals. There are no differences in the number of puncta at P30, however these are significantly reduced by P90 (each point represents a field analyzed, 5 images analyzed per biological replicate, n= 3 biological replicates. p = 0.0098 by One-way anova with Tukey correction, error bars indicate SEM 72.93 +/- 4.62). **(E)** Accelerating rotarod performance test of *Ttbk2^PCP2^* and littermate controls at aging from P30 to P90. *Ttbk2^PCP2^* animals have a shorter latency to fall time at P30 and P90. **(F)** Steady speed rotarod performance test of *Ttbk2^PCP2^* and littermate controls aging from P30 to P90. At P30 *Ttbk2^PCP2^* animals do not have a shorter latency to fall time compared to controls on the steady speed rotarod. However, by P90 there is a drastic reduction latency to fall time for *Ttbk2^PCP2^* animals compared to controls, indicative of impaired motor ability with age (n=6 biological replicates for Control, n=4 biological replicates for *Ttbk2^PCP2^*).

We then tested P30 and P90 *Ttbk2^PCP2^* animals using the rotarod performance test as an assay for motor coordination, and did not observe significant changes in P30 animals. However, by P90, the *Ttbk2^PCP2^* animals consistently exhibited reduced latency to fall on both the accelerating rotarod as well as the steady state rotarod performance tests (Fig. 3E, 3F). These data show that loss of *Ttbk2* from PCs causes neurodegenerative phenotypes to arise over time. Overall, these phenotypes are less severe than those of the *Ttbk2^c.mut^* animals, implying that other cell types may also contribute to the loss of synaptic puncta. Taken together, this data reveals that *Ttbk2* plays a critical role in maintaining the architecture and connectivity of the cerebellum. Within three months following genetic loss of *Ttbk2*, we observe degenerative changes to several cell types within the cerebellum that comprise an important neural circuit: synaptic connections are reduced between the PCs and the neurons of the ION. The cell bodies of the ION neurons also shrink significantly during this time and the PCs exhibit some thinning of their large, elaborate dendrites. BGs also exhibit alterations in their morphology upon loss of TTBK2. These changes are accompanied by motor coordination defects in these animals.

### Loss of *Ttbk2* ultimately leads to loss of Purkinje Cells

In the first 3 months following TMX injections, the phenotypes exhibited by the *Ttbk2^cmut^* mice consisted mainly of altered synaptic connectivity between PC and ION climbing fibers, and accompanying deficits in motor coordination. However, when we assessed the cerebellar phenotypes of animals at 6 months of age (5 months following TMX injection), we found regions throughout the cerebellar folia where the soma of the PCs were absent, often appearing as gaps in the molecular layer upon staining with Calbindin to label PCs (Figure 4A, 4B). We counted the number of PC soma within a defined region of the primary fissure and found that these are reduced in 6 month old *Ttbk2^c.mut^* mice compared to littermate controls of the same age, as well as compared to 4 month old *Ttbk2^c.mut^* mice (Figure 4B. 18.5 +/- 0.29 PCs per 500 μm for 4 month Control vs. 18.42 PCs +/- 0.34 for 4 month *Ttbk2^c.mut^*. 18.67 PCs +/- 0.43 for 6 month Control vs. 11.92 PCs +/- 0.74 for 6 month *Ttbk2^c.mut^*). These results indicate that prolonged loss of *Ttbk2* leads to neurodegeneration and loss of PCs throughout the PC layer of the adult cerebellum.

**Figure 4.**
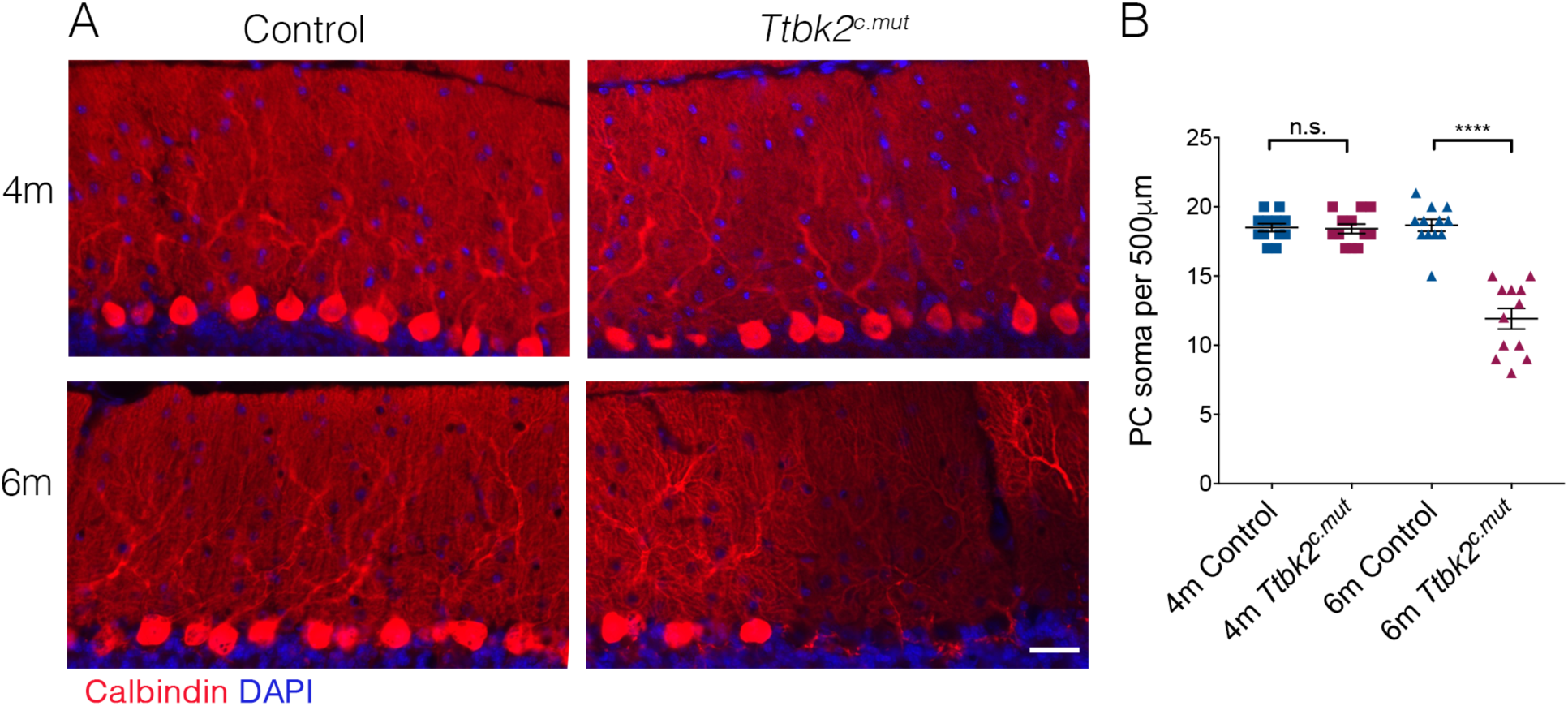
Aged *Ttbk2^c.mut^* animals lose Purkinje cells. **(A)** Representative images showing folia of 4m old *Ttbk2^c.mut^* (top) and 6m old *Ttbk2^c.mut^* animals (bottom) with respective littermate controls. Cerebellum tissue is stained for Calbindin to show PC. 6mo *Ttbk2^c.mut^* have large stretches of folia missing Calbindin+ PC soma compared to 4m *Ttbk2^c.mut^*. Scale bar 50μm. **(B)** Quantification of the loss of PC soma along 500μm stretch of the primary fissure. (n=36 measurements across 3 biological replicates. p<0.0001 by student’s unpaired t-test, error bars indicate SEM 6.75 +/- 0.86).

### *Ttbk2^c.mut^* animals lack neuronal primary cilia

Our prior work demonstrated that mutations associated with SCA11, which result in the expression of a truncated protein, interfere with the function of full-length TTBK2. In particular, these mutations dominantly interfere with cilia formation in embryos and cultured cells (Bowie et al., 2018). Throughout the adult cerebellum and other regions of the hindbrain, neurons as well as glia possess primary cilia (Figure 5A-5C). Within 20 days following administration of TMX to induce recombination (P45), the number of ciliated cells in the cerebellum declined dramatically in *Ttbk2^c.mut^* mice: from a mean of 22.46 cilia per 32mm^2^ field +/- 0.7626 in control animals to 2.36 cilia per 32mm^2^ field +/- 0.3103 in *Ttbk2^c.mut^* animals (Figure 5D, 5E). This loss of cilia was observed throughout the cerebellum, brain stem, and other areas of the brain which we examined such as the hippocampus and the cortex (Figure S3A and S3B). This loss of cilia precedes the behavioral and cellular changes we have identified throughout the cerebellum. Consistent with other conditional mutants where cilia are globally removed in adulthood (Davenport et al., 2007) 4 month old *Ttbk2^c.mut^* mice exhibit obesity (Figure S4A, A’ and Figure S4B: 32.29 g +/- 1.86 for Control vs. 46.33 g +/- 2.04 for *Ttbk2^c.mut^*) as well as cystic kidneys (Figure S4C). Taken together this data illustrates that *Ttbk2* is required for neuronal cilia throughout the adult mouse brain, which is in line with our current data showing that *Ttbk2* is required for ciliogensis and cilium stability.

**Figure 5.**
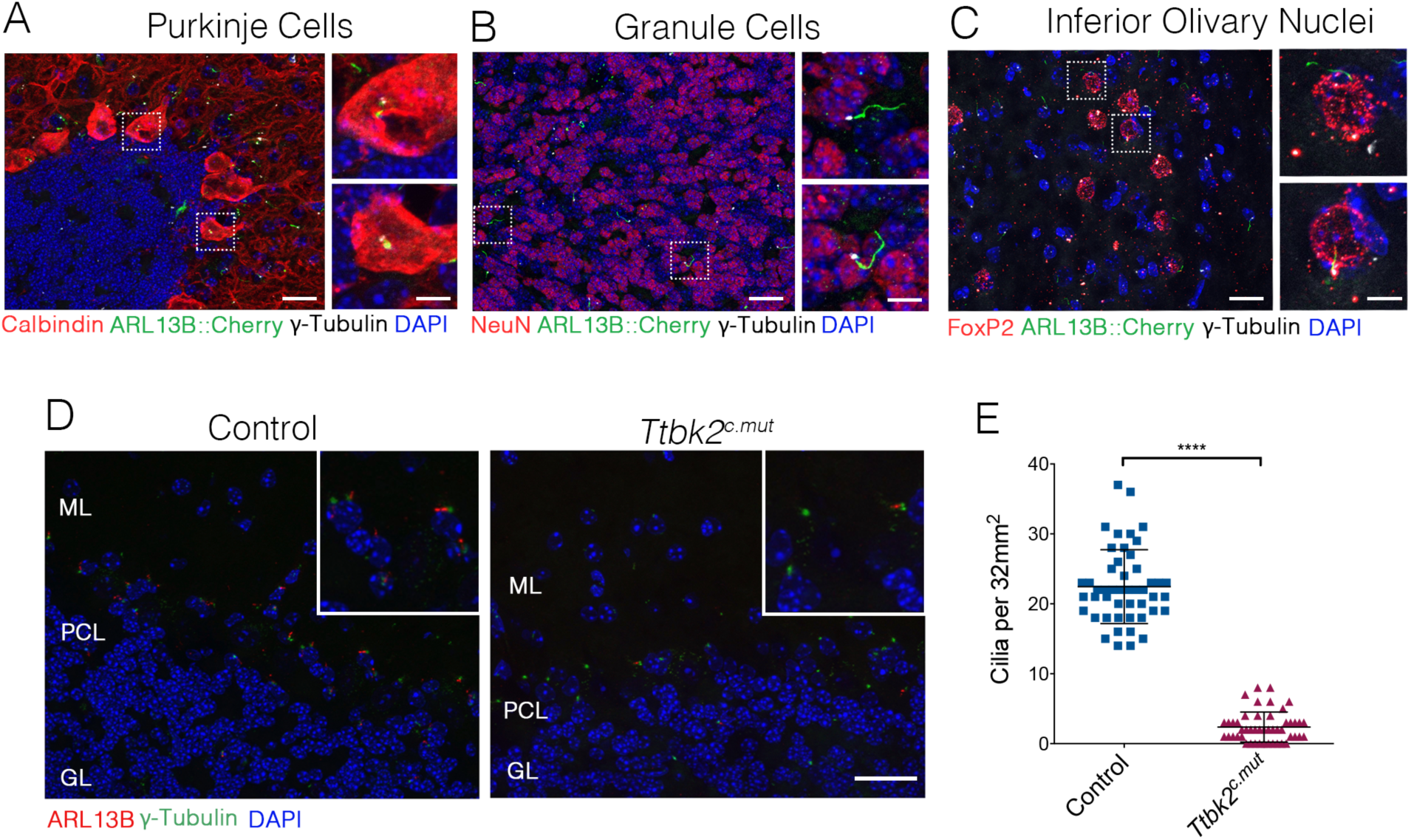
*Ttbk2* is critical for primary cilia stability on neurons. **(A-C)** Representative images of cilia on specific cell types throughout parts of the cerebellum and medulla. Sections from a mouse expressing the knock in allele for ARL13B tagged with cherry were immunostained for γ- Tubulin to label centrosomes (magenta), and various cell specific markers such as Calbindin to label Purkinje Cells (red, **A**), NeuN to label granule neurons (red, **B**), and FoxP2 to label neurons within the Inferior Olivary Nucleus (red,**C**). Insets show boxed areas. Scale bar = 50μm, 10μm for insets. **(D)** Representative images illustrating cilia loss in the cerebellum twenty days after tamoxifen treatment. Sections were immunostained for ARL13B to label cilia (red) and γ- Tubulin to label centrosomes (green). For quantification purposes images were taken at the nexus between the molecular layer (ML) and granule layer (GL) with the Purkinje cell layer (PCL) in the middle of the imaging field where there is an abundance of cilia. Scale bar =50μm. **(E)** Quantification of cilia loss after TMX treatment (n = 36 images counted, 3 biological replicates, p<0.0001 student’s unpaired t-test, error bars indicate SEM 20.08 +/- 0.8233).

### Loss of *Ift88* recapitulates *Ttbk2^c.mut^* phenotypes

*Ttbk2* is essential both for the initiation of cilium assembly as well as for the structure and stability of cilia (Bowie et al., 2018; Goetz et al., 2012). Consistent with this, we have found that primary cilia are lost from cells within the cerebellum of *Ttbk2^cmut^* mice (Figure 5) prior to the onset of the phenotypes we observe in neuronal connectivity and survival. Given this critical link between TTBK2 and primary cilia in all cell types examined both during development as well as in adult tissues, we next tested whether loss or dysfunction of cilia function via a different genetic mechanism causes convergent phenotypes to those of the *Ttbk2^cmut^* mice. For these studies we turned to conditional mutants of another key ciliary protein, Intraflagellar Transport Protein 88 (IFT88). IFT88 is a component of the IFTB particle required for assembly of the ciliary axoneme as well as anterograde trafficking within the cilium (Pazour et al., 2000). Our previous work shows that IFT88 functions downstream of TTBK2 in cilium initiation (Goetz et al., 2012), with TTBK2 being required IFT recruitment. A hypomorphic allele of *Ift88* in the mouse, *Tg737^orpk^* (Pazour et al., 2000) causes cystic kidneys and other phenotypes reminiscent of human ciliopathies (Reiter and Leroux, 2017). In the brain, IFT88 is important for cilia structure in the hippocampus and cortex, and when knocked out in these specific neuron populations results in memory deficits (Berbari et al., 2014). Additionally, *Ift88* null mutants exhibit nearly identical embryonic phenotypes to those of *Ttbk2* null mutants (Murcia et al., 2000). When we knocked out *Ift88* using the same methods described for *Ttbk2^c.mut^* animals, we see that at 3 months post TMX treatment, more cilia remain throughout the cerebellum in *Ift88^c.mut^* cerebella compared to the *Ttbk2^c.mut^* animals with the same treatment (Figure S5A, S5C; 17.31 cilia per 32mm^2^ field +/- 0.65 for Control vs. 12.13 cilia per 32mm^2^ field +/- 0.50 for *Ift88^c.mut^*). However, the cilia that remain in *Ift88^c.mut^* animals are shorter in length than controls (Figure S5C; 2.31μm +/- 0.10 for control vs. 1.70μm +/- 0.08 for *Ift88^c.mut^*). Western blot analysis confirmed a loss of IFT88 protein in brain lysates from four month old *Ift88^c.mut^* cerebellar lysate (Figure S5B). This analysis confirms that *Ift88^c.mut^* mice lose IFT88 protein, and this results in reduced numbers of shortened cilia throughout the cerebellum.

Further analysis of cilia status in *Ift88^c.mut^* mice revealed changes in localization of key signaling molecules associated with neuronal cilia. Neuronal cilia express a number of membrane proteins important for various signaling cascades, including Adenylate Cyclase 3 (AC3)(Guadiana et al., 2016). We observed that within the cerebellum, there is a distinct population of cilia which are AC3+/ARL13B- (Figure S6A, arrowhead), while some cilia are double positive for AC3+/ARL13B+ (Figure S6B), and another population has only ARL13+ cilia. In our *Ift88^c.mut^* animals, we see various changes to these populations, specific to their AC3 signature. The population of AC+ only cilia are nearly completely lost in *Ift88^c.mut^* animals (Figure S6A, S6C: 3.42 cilia per 32mm^2^ field +/- 0.32 in Control vs. 0.22 cilia per 32mm^2^ field +/- 0.09 in *Ift88^c.mut^*), reminiscent of the drastic loss of ARL13B+ cilia in the *Ttbk2^c.mut^* animals. Additionally, there is a strong reduction in AC3+/ARL13B+ double positive cilia in *Ift88^c.mut^* (Figure S6B, S6D: 4.06 cilia per 32mm^2^ field +/- 0.24 in Control vs. 1.94 cilia per 32mm^2^ field +/- 0.20 in *Ift88^c.mut^*). This analysis shows that IFT88 is required for the localization of specific signaling molecules such as AC3 to neuronal primary cilia throughout the cerebellum.

We then examined cerebellar structure and circuitry in *Ift88^c.mut^* animals. Similar to our findings in *Ttbk2^c.mut^* animals, no changes to PC number were evident at 3 months post TMX treatment. Molecular layer thickness was reduced in *Ift88^c.mut^* animals (Figure 6A, 6B. 182.3μm +/- 2.5 in Control vs. 169.3μm +/- 3.04 in *Ift88^c.mut^*), though this change was less severe than what we observed in *Ttbk2^c.mut^* mice. Similarly, VGLUT2 puncta were reduced in *Ift88^c.mut^* compared to controls (Figure 6A, 6C. 515.7 puncta +/- 20.58 in control vs. 395.6 puncta +/- 13.7 in *Ift88^c.mut^*). We next tested *Ift88^c.mut^* animals on the rotarod performance test to uncover any motor coordination deficits, given that these mice exhibit similar cellular changes to those of the *Ttbk2^cmut^* animals. These tests revealed that *Ift88^c.mut^* animals also have a shorter latency to fall time on the steady speed rotarod performance test, but not on the accelerating rotarod performance test. This is in contrast to *Ttbk2^c.mut^* animals which have a shorter latency to fall time on both accelerating and steady speed rotarod performance tests. Taken together, this data suggests that IFT88 is important for cilia structure and function throughout the cerebellum, though to a different degree than that of TTBK2. Unlike where we see an almost complete loss of cilia throughout the cerebellum in *Ttbk2^c.mut^* animals, *Ift88^c.mut^* have more cilia remaining after knockout, and these cilia that remain are overall shorter than controls, indicating to us that trafficking along the ciliary axoneme is impaired.

**Figure 6.**
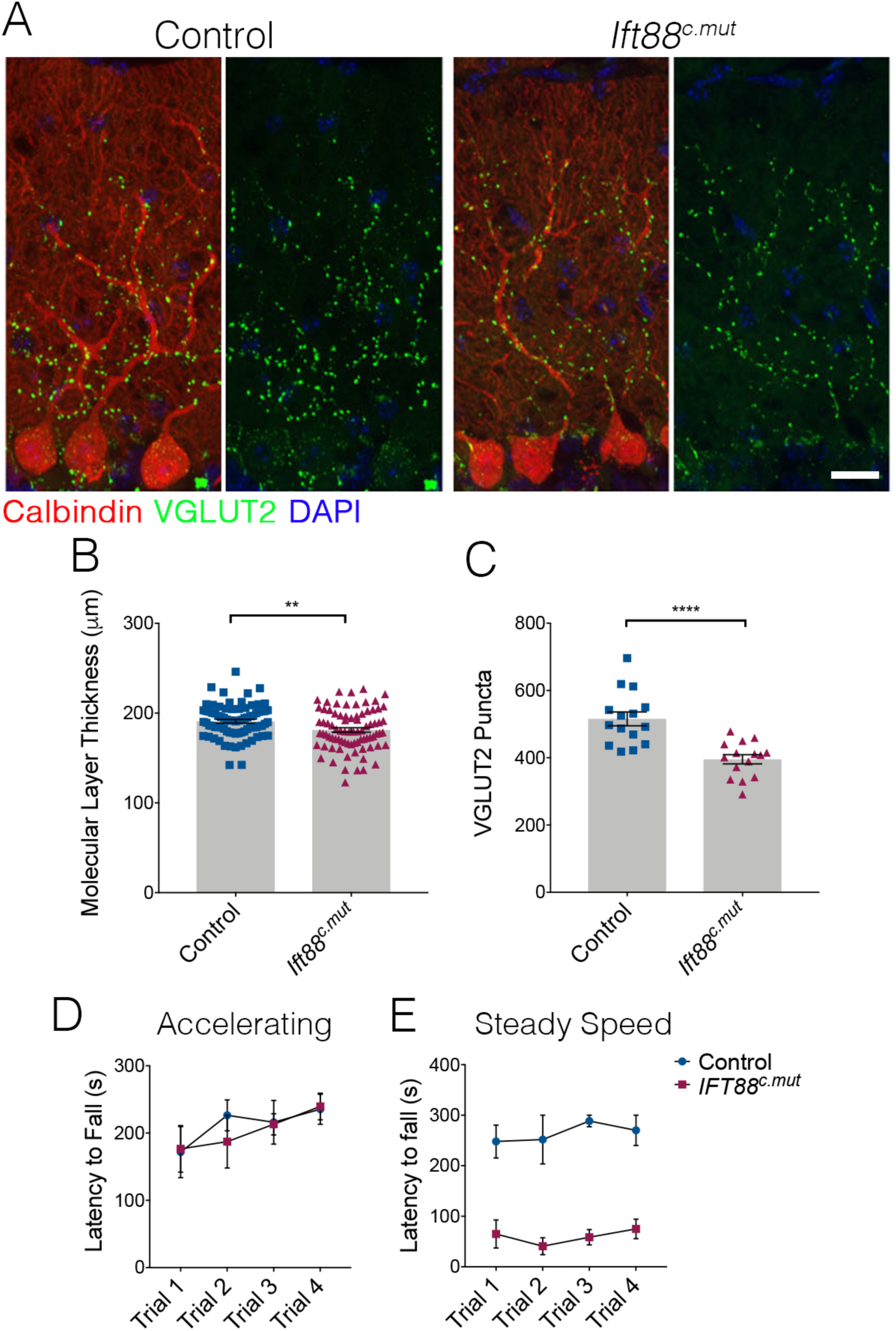
Loss of *Ift88* recapitulates neurodegenerative phenotypes of *Ttbk2^c.mut^* animals. **(A)** Cerebellar tissue from control and *Ift88^c.mut^* mice at three months after loss of *Ift88*, immunostained for Calbindin to label Purkinje cells (red) and VGLUT2 to show climbing fiber synapses (green). *Ift88^c.mut^* animals show a reduction in VGLUT2 positive synapses throughout the cerebellum three months after loss of IFT88. Scale bar = 50μm. **(B)** Molecular layer length quantification of *Ift88^c.mut^* animals compared to littermate controls. Each point represents one measurement, >75 measurements taken per genotype. n=3 biological replicates. p=0.0037 by unpaired student’s t-test, error bars indicate SEM 9.60 +/- 3.26. **(C)** Quantification of loss of VGLUT2 synapses along PC dendrites in *Ift88^c.mut^* animals. Each point represents a field analyzed, 5 images analyzed per biological replicate, n= 3 biological replicates. p<0.0001 by unpaired student’s t-test, error bars indicate SEM 120.1 +/- 24.72. **(D, E)** Accelerating and Steady Speed Rotarod Performance Test between *Ift88^c.mut^* and littermate controls. *Ift88^c.mut^* animals have a shorter latency to fall time on the Steady Speed Rotarod test, but did not have any significant difference on the Accelerating Rotarod test (n=6 biological replicates for Control, n=4 biological replicates for *Ift88^c.mut^*).

Because the phenotypes we observed in *Ift88^c.mut^* at 3 months post TMX were somewhat milder than the changes we observed in *Ttbk2^c.mut^* animals of the same age, we next asked if we would see loss of PCs in aged *Ift88^c.mut^*. In six month old *Ttbk2^c.mut^* we begin to see loss of PCs throughout the cerebellum, which show up as large gaps in the folia (Figure 4). Similar to these results, 6 month old *Ift88^c.mut^* animals show gaps throughout the PC layer, and have reduced numbers of PC soma, similar to the *Ttbk2^cmut^* mice (Figure 7A, 7B; 17.5 +/- 0.44 per 500μm in 4mo control vs. 16.92 PC soma +/- 0.31 in 4mo Ift88*^c.mut^*. 17.17 PC soma +/- 0.55 in 6mo control vs. 12.67 PC soma +/- 0.43 in 6mo *Ift88^c.mut^*). Coupled with these findings, the molecular layer thickness is further reduced in 6mo *Ift88^c.mut^* animals (Figure 7C, 7D, 7E; 173.9μm +/- 2.28 in 6mo control vs. 158.0μm +/- 1.63 in 6mo *Ift88^c.mut^*), as well as VGLUT2 puncta counts were diminished in 6 month old *Ift88^c.mut^* t (Figure 7C, 7D, 7F; 645.9 puncta +/- 26.83 in 6mo control vs. 461.4 puncta +/- 25.42 in 6mo *Ift88^c.mut^*). These changes confirm that loss of *Ift88* throughout the adult cerebellum causes neurodegeneration to occur, and PC loss to begin at six months of age.

**Figure 7.**
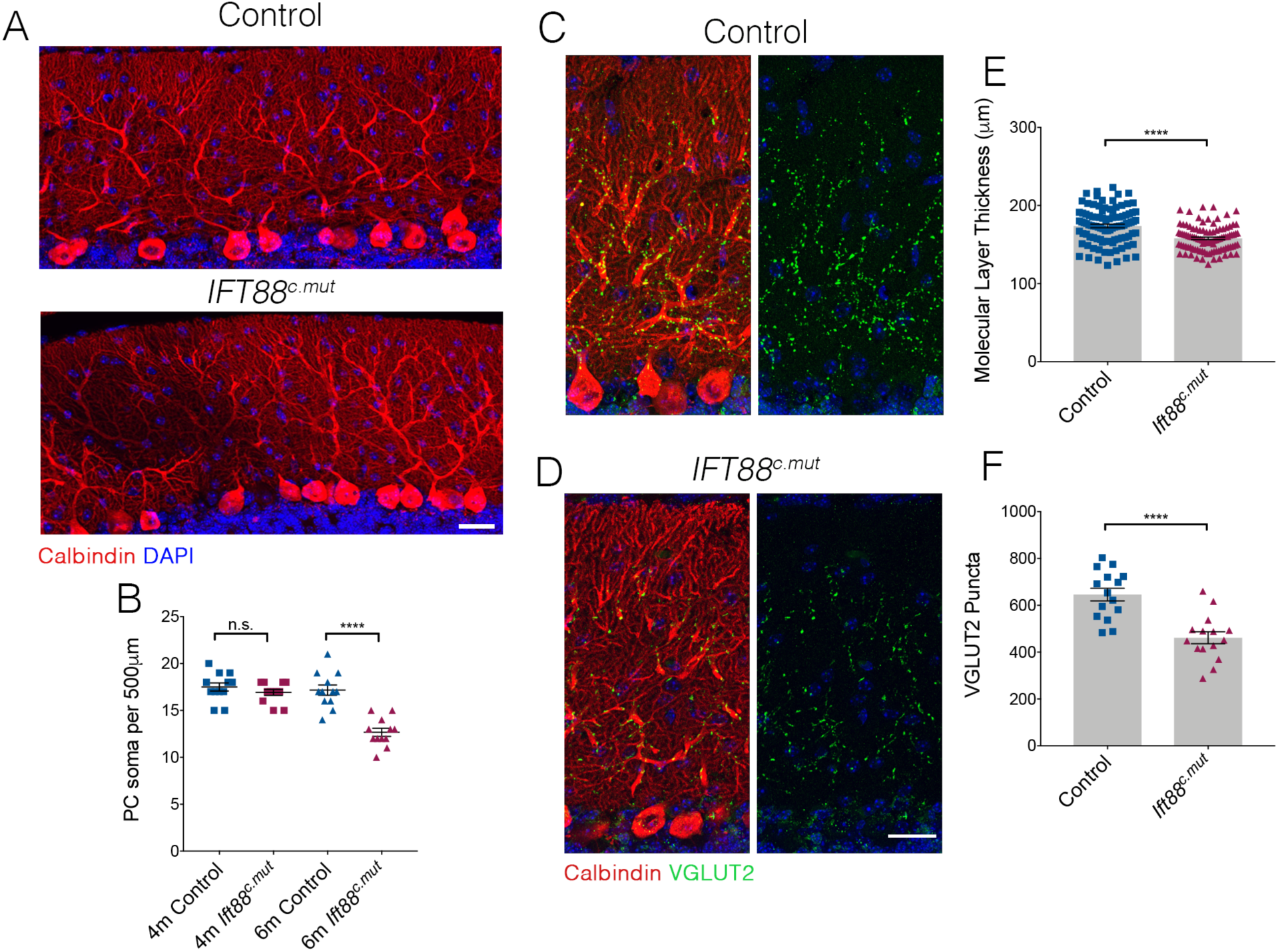
Loss of PCs occurs in *Ift88^c.mut^* mice by 6 months of age. **(A)** Cerebellar folia from Control and *Ift88^c.mut^* at 6 months of age, five months after loss of *Ift88*, stained for Calbindin (red) and dapi (blue). Six month old *Ift88^c.mut^* mice begin to show PC gaps throughout cerebellar folia indicating loss of PC. Scale bar=50μm. **(B)** Quantification of number of PC soma per 500μm stretch of folia on the primary fissure between Control and *Ift88^c.mut^* at 4 months and 6 months of age. Six month old *Ift88^c.mut^* show a reduced number of PC soma throughout the cerebellum (each point represents one measurement, twelve measurements were made per condition. n=3 biological replicates per condition. p<0.0001 by one-way ANOVA between all conditions, error bars indicate SEM). **(C, D)** 6 month old cerebellar tissue from Control and *Ift88^c.mut^* mice immunostained for Calbindin to label Purkinje cells (red) and VGLUT2 to show climbing fiber synapses (green) and dapi (blue). *Ift88^c.mut^* animals show a reduction in VGLUT2 positive synapses throughout the cerebellum. Scale bar = 50μm. **(E)** Quantification of molecular layer thickness between 6 month old Control and *Ift88^c.mut^* animals. *Ift88^c.mut^* have shorter folia compared to littermate controls (each point represents one measurement, >75 measurements taken per genotype. n=3 biological replicates. p<0.0001 by unpaired student’s t- test, error bars indicate SEM -15.91+/- 2.81). **(F)** Quantification of loss of VGLUT2 synapses along PC dendrites in 6 month *Ift88^c.mut^* animals. *Ift88^c.mut^* a loss similar to the loss seen in 4 month old *Ift88^c.mut^* animals (each point represents a field analyzed, 5 images analyzed per biological replicate, n= 3 biological replicates. p<0.0001 by unpaired student’s t-test, error bars indicate SEM -184.5 +/- 36.96).

## DISCUSSION

In this work, we have tested the hypothesis that SCA11 pathology results from the requirements for TTBK2 in cilium assembly and stability. We show that TTBK2 is essential for maintaining the connectivity and viability of PCs in the adult cerebellum. These phenotypes are similar to those reported for mouse models of other subtypes of SCA as well as consistent with many aspects of SCA11 in human patients, suggesting that the *Ttbk2^cmut^* mice model the human condition. We further demonstrate that mice conditionally lacking the ciliary protein IFT88 in adult tissues exhibit highly similar neurodegenerative phenotypes to those that we observe in the *Ttbk2^cmut^* mice, including loss of excitatory synapses from the climbing fibers and eventual loss of PCs. The high degree of convergence of these phenotypes suggests that the degenerative phenotypes that occur in the *Ttbk2^cmut^* mice are driven primarily by the requirement for TTBK2 in mediating cilium assembly, and point to a critical role for these organelles in maintaining neuronal function during adulthood.

Cilia and ciliary signals play a variety of important roles during embryonic and postnatal development of the brain and central nervous system. Cilia are linked to processes including the expansion and patterning of neural progenitors(Guemez-Gamboa et al., 2014), the migration and laminar placement of interneurons(Higginbotham et al., 2012), and in the establishment of neuronal morphology(Guadiana et al., 2016; Sarkisian and Guadiana, 2015). Consistent with the varied roles of cilia in neural development, an array of neurological deficits are among the most common hallmarks of ciliopathies(Lee and Gleeson, 2010; Youn and Han, 2018), highlighting the importance of these organelles in human health. In addition to their critical developmental functions, mounting evidence supports an important role for ciliary signaling in tissue regeneration and homeostasis in many adult organs, including the kidneys(Davenport et al., 2007), skin (Croyle et al., 2011), skeletal muscle(Kopinke et al., 2017) and bone(Moore et al., 2018).

Within the adult CNS, dysfunction of ciliary trafficking is linked to retinal degeneration(Wheway et al., 2014), and conditional loss of the ciliary protein ARL13B from mouse striatal interneurons both during their development as well as in the mature brain, results in changes in their morphology and connectivity(Guo et al., 2017). Our work extends these findings and provides additional strong evidence that primary cilia and signals mediated by these organelles are important to maintain the morphology of neurons as well as their connections. In addition, we have shown that a specific type of neuron within the brain, cerebellar PCs, requires a functional primary cilium for their survival. Degeneration of a specific type of neuron, the photoreceptors of the retina, is well known to occur under conditions where cilia are dysfunctional- both in many human ciliopathies as well as in mouse models of these disorders(Braun and Hildebrandt, 2017; Bujakowska et al., 2017). In the case of photoreceptors, degeneration occurs as trafficking within the outer segment fails, resulting in accumulation of rhodopsin within the cell body and leading to photoreceptor cell death through mechanisms that are not completely understood(Seo and Datta, 2017). In the case of PCs, loss of cilia leads to changes in Ca++ signaling concomitant with reduced numbers of excitatory synapses from the climbing fibers of the ION neurons.

While the precise nature of the ciliary signals that maintain the connectivity and viability of neurons remains unknown, there are a number of candidates. Many different GPCRs and associated signaling cascades and second messengers have been shown to concentrate in primary cilia or to be enriched predominantly at the primary cilium(Mykytyn and Askwith, 2017). In particular cAMP and Ca++ are highly concentrated within the cilium (Moore et al., 2016). Misregulation of these concentrations through either ciliary loss or dysfunction could therefore be expected to result in perturbed signaling outputs to the cell bodies of these neurons. Additionally, primary cilia play a well-known role as essential mediators of Hedgehog (HH) signaling. Sonic Hedgehog (SHH) is secreted by PCs both during development as well into adulthood(Lewis et al., 2004; Traiffort et al., 1998), although the precise role and functional significance of SHH within the adult cerebellum is largely unclear. Ultimately, it will be important to investigate the regulation and molecular composition of neuronal cilia in a comprehensive and unbiased manner. Our data show that conditional mutants of *Ttbk2* and *Ift88* have similar phenotypes with respect to loss of excitatory synapses to PCs from the climbing fibers, motor coordination deficits, and eventually loss of PCs. This evidence suggests that these defects are the result of ciliary loss or dysfunction. We note however, that the ciliary phenotypes that result from loss of *Ttbk2* differ from those observed in the *Ift88* conditional mutants in the context of the adult brain. *Ttbk2* conditional mutant mice rapidly lose cilia following administration with TMX, with nearly all cilia within the cerebellum being absent within 14 days. In contrast, the numbers of cilia in the *Ift88* conditional mutants are only slightly reduced 3 months following TMX. However, these cilia exhibit significant abnormalities, including reduced length, and a near-complete loss of adenylate cyclase 3 (AC3) from the remaining cilia. This suggests strongly that the degenerative phenotypes observed in both conditional mutants are driven by the loss of a specific ciliary signal.

These observations also have interesting implications for the regulation of cilium assembly and stability both generally and in post-mitotic adult cells and neurons in particular. We find for example, that IFT88 protein is lost in *Ift88* conditional mutants as expected. However the cilia persist on adult neurons within the brain in the absence of IFT88, suggesting that fully functional IFT is not required for these cilia. In contrast, in the absence of TTBK2, cilia are rapidly lost. This suggests that TTBK2 is playing a more central role in maintaining the stability of cilia in this context than IFT machinery. The exact mechanisms by which TTBK2 regulates the stability of cilia and the degree to which this role may be specifically important in neurons will be the subject of future investigations within our lab. In particular, the dynamics of cilium assembly and disassembly in post-mitotic cells *in vivo* have not been characterized. For example, a recent study found that the proteins that comprise the basal body of adult neurons in the mouse are very long-lived while those of the ciliary axoneme turn over more quickly(Drigo et al., 2019). This might imply that the cilia of these adult neurons turn over at some interval, or simply that their protein components are replaced- a topic that merits further investigation.

In addition to being required for the biogenesis of cilia, TTBK2 also localizes to the + tips of microtubules, mediated through its interaction the + end binding protein EB1 (Jiang et al., 2012). TTBK2 has also been shown to phosphorylate β-Tubulin as well as microtubule associated proteins TAU and MAP2 through in vitro assays (Takahashi et al., 1995; Tomizawa et al., 2001), pointing to roles for TTBK2 in the regulation of the microtubule cytoskeleton beyond the cilium. In addition, TTBK2 and the closely related kinase TTBK1 both phosphorylate SV2A *in vitro*, a component of synaptic vesicles important for the retrieval of the membrane trafficking protein Synaptotagmin 1 during the endocytosis of synaptic vesicles (Zhang et al., 2015). While our data showing that the *Ttbk2* mutant phenotypes strongly overlap with those of other ciliary genes, such as *Ift88*, we can not exclude the possibility that other roles of TTBK2 specifically within the brain also contribute to the degenerative phenotypes. Importantly, these two possibilities are not mutually exclusive, and indeed, one exciting possibility is that TTBK2 is important for relaying signals from the cilium to the neuronal cell body.

## CONCLUSIONS

In this work, we present evidence that loss or impaired function of TTBK2 within the brain results in degeneration of PCs due largely to the requirement for TTBK2 in mediating the assembly and stability of primary cilia. This points to ciliary dysfunction as being a major mechanism underlying the pathology of SCA11, which is caused by truncating mutations to TTBK2 (Houlden et al., 2007) that act as dominant negatives (Bowie et al., 2018). In addition, our work raises the possibility that cilia play an important, largely unappreciated role in maintaining neuronal connectivity within the brain, and may also be required for the viability of some types of neurons.

From a clinical perspective, our findings suggest that neurodegeneration, in addition to other neurological impairments with a developmental origin, may emerge in some patients with ciliopathies such as Joubert and Bardet Biedl syndromes, particularly as patients age.

## MATERIALS AND METHODS

### Ethics Statement

The use and care of mice as described in this study was approved by the Institutional Animal Care and Use Committees of Duke University (Approval Number A218-17-09). All animal studies were performed in compliance with internationally accepted standards.

### Mouse Strains

*Ttbk2^c.mut^* mice were produced by crossing *Ttbk2^tm1a(EUCOMM)Hmgu^* mice to ACTB:FLPe (Jax stock #003800). The following mice were purchased from Jackson Laboratories: *Ift88^flox^* (stock #022409), UBC-CreER (stock #007001), and PCP2-Cre (Jax stock #010536, gift from Dr. Court Hull at Duke University).

### Genotyping

PCR genotyping was performed on all mice before experiments to confirm presence of floxed alleles and Cre. *Ttbk2*-floxed allele, primers used: 5’ ATACGGTTGAGATTCTTCTCCA, 3’ AGGCTGTACTGTAACTCACAAT (WT band 978bp, floxed band 1241bp). *Ift88*-floxed allele, primers used: 5’ GCCTCCTGTTTCTTGACAACAGTG, 3’ GGTCCTAACAAGTAAGCCCAGTGTT (WT band 350bp, floxed band 370bp). Universal Cre (UBC-CreER, PCP2-Cre), primers used: 5’ GATCTCCGGTATTGAAACTCCAGC, 3’ GCTAAACATGCTTCATCGTCGG (transgene band 650bp).

### Tamoxifen preparation and injection

Tamoxifen powder (Sigma T5648) was dissolved in corn oil (Sigma C8267) to a desired concentration of 20mg/mL. Mice were given five consecutive 100μL intraperitoneal injections of 20mg/mL tamoxifen starting at P21. Control mice were given corn oil vehicle only.

### Mouse Dissections

To harvest tissues from adult mice, animals were deeply anesthetized with 12.5mg/mL Avertin and transcardially perfused with 10mL of Phosphate Buffered Saline (PBS) followed by 20mL of 4% Paraformaldehyde (PFA). Whole brains were dissected out and left to incubate for 24hrs in 4% PFA at 4°C. For cryosectioning, tissue was cryoprotected in 30% sucrose overnight and embedded in Tissue Freezing Medium (General Data TFM-5). Cerebella were then cut sagittally down the middle, and embedded in Tissue Freezing Medium (General Data TFM-5). Tissue was sectioned at 20-30μm thickness on a Leica Cyrostat (model CM3050S).

### Western Blotting

Western blot methods were done as previously described(Bouskila et al., 2011) (Bowie et al., 2018). Briefly, for tissue which was being used to quantify levels of TTBK2, a buffer containing 50 mM Tris/HCl, pH 7.5, 1 mM EGTA, 1 mM EDTA, 1 mM sodium orthovanadate, 10 mM sodium-2-glycerophosphate, 50 mM sodium fluoride, 5 mM sodium pyrophosphate, 0.27 M sucrose, 1 mM benzamidine and 2 mM PMSF, supplemented with 0.5% NP-40 and 150 mM NaCl was used. For other tissue samples a buffer containing 10mM Tris/Cl pH7.5, 150mM NaCl, 0.5mM EDTA, 1% Triton, 1mM protease inhibitors (Sigma #11836170001) and 25mM β- glycerol phosphate (Sigma 50020) was used. Total protein concentration was determined using a BSA Protein Assay Kit (Thermo Fisher #23227). For western blots, 15μg of protein lysate was used for detection.

### Cilia quantification

All quantification of cerebellar tissue was done using ImageJ software. Images taken for quantification of cilia abundance were 10μm z-stacks taken at 63x in four distinct folia regions of the cerebellum, two rostral and two caudal. The Purkinje cell layer was placed into the middle of the image with equal distance above and below for quantification. The outer edge of folia 2, the internal zone between folia 3 and 4, the tip of folia 6, and the outer edge of folia 9 on a sagittal section were imaged for cilia quantification. Per replicate, four sections were scored each and three biological replicates were included in all quantifications. These cilia were the same population taken for cilia length measurements as well.

### Molecular layer thickness and VGLUT2 puncta quantification

For the molecular layer thickness, images were taken at 20x along the entirety of the primary fissure. A line was drawn from the bottom of the molecular layer to the pial surface, and a measurement was recorded. For this same line, the top of the line measurement was then brought down to the top of the VGLUT2 synapse area, and a measurement recorded. Only the caudal side of the folia was measured for consistency. For the VGLUT2 puncta analysis, the “Analyze Particles” function in ImageJ was used. Each image for the VGLUT2 puncta analysis was taken at 63x on the caudal side of the primary fissure, a 10μm z-stack was made, and the image quantified. For the quantification, each stack was made into a black and white image, where the VGLUT2 puncta were black against a white background. Thresholding was performed, and the Analyze Particle function used. These measurements were routinely tested against user ROI counting to confirm accuracy. Images used for cilia counts were taken at 63x. A 10μm z-stack was taken at 4 locations throughout the cerebellum. The image was taken with the PCL in the middle of the image for consistency. Four cerebellum slices were imaged per biological replicate..

### Inferior Olivary Nuclei Quantification

For the area measurements of the ION nuclei, the ION was identified by cells that were positive for both NeuN and Calbindin as well as in the part of the ventral medulla in which these cells reside. Images used for the NeuN area analysis were taken at 20x. A 10μm z-stack image was made, and using the line tool, outlines were carefully drawn around the NeuN positive neuron and the measurement recorded. Per replicate, over 150 cells were measured and three biological replicates were included in the quantification.

### Glial Fiber Quantification

Glial fibers were assessed as previously described (Furrer et al., 2011). Briefly, a 100μm horizontal line was drawn 50μm below the pial surface of the primary fissure folia. Glial fibers which crossed this 100μm were scored. Per replicate, 36 measurements were made and three biological replicates were included in the quantification.

### Immunostaining

The following antibodies and dilutions were used in this study: mouse anti-Arl13b (NeuroMabs N295B/66, 1:500), rabbit anti-Arl13b (gift from Tamara Caspary), mouse anti-gamma-Tubulin (Sigma T6557, 1:1000), rabbit anti-Calbindin D28K (Cell Signaling Technologies 13176S, 1:250), guinea pig anti-Calbindin D28K (Synaptic Systems 214-004, 1:200), guinea pig anti- VGLUT2 (EMD Millipore AB2251, 1:2500), rabbit anti-NeuN (Abcam ab177487, 1:1000), DAPI (Sigma D9542, 1x), rabbit anti-AC3 (Santa Cruz SC-588, 1:10 - discontinued), chicken anti-GFAP (EMD Millipore AB5541, 1:500), rabbit anti-IP3 (Abcam, ab108517, 1:200).

For Immunostaining cerebellar tissue, after thawing from the freezer, sections were rinsed in 1xPBS to remove OCT product. Following rinse, sections were permeabilized in 0.2% PBS-T (PBS + 0.2% Triton X-100) for 10 minutes, and then rinsed 3x5min in PBS before blocking step. Blocking solution contained 5% Serum, 1% BSA made up in 0.1% PBS-T, and sections were incubated at room temperature in blocking solution for 1 hour. Primary antibodies were used at indicated dilutions and incubated at 4°C overnight. Following primary antibody incubation, slides were rinsed 3x5min in 1xPBS and secondary antibodies were used to detect epitopes. All secondary antibodies were supplied from Life Technologies. Secondary antibodies incubated for 1-3hr at room temperature. Following secondary antibody incubation, slides were rinsed 3x5min in 1xPBS and mounted with either ProLong Gold antifade reagent (Invitrogen P23930).

### Behavioral Testing

Rotarod Performance Test was completed with help from the Duke University Mouse Behavioral and Neuroendocrine Core Facility. Testers were blind to mouse genotype before beginning any experiments. The accelerating rotarod testing was performed the day before steady state rotarod testing. The speed determined for the steady speed testing was calculated based off results from the accelerating test (4-40RPM) the day prior. In all tests, a trial was stopped after 300 seconds maximum time had elapsed for mice that did not fall off the rotarod during testing. Mice were aborted from the trial run if they held onto the Rotarod for three full rotations. Mice were given 30 minutes between trials to rest, and four trials were completed per test.

## ACKNOWLEDGEMENTS

We are grateful to William Wetsel, Romona Rodriguiz, and other staff of the Duke University Mouse Behavior and Neuroendocrine Core Facility for assistance with behavioral testing. We thank members of the Goetz lab for helpful comments on the manuscript. This work was supported by a grants from NIH/NICHD (R00HD076444) and the National Ataxia Foundation (Young Investigator Award in SCA) to SCG.

---

**Figure S1.**
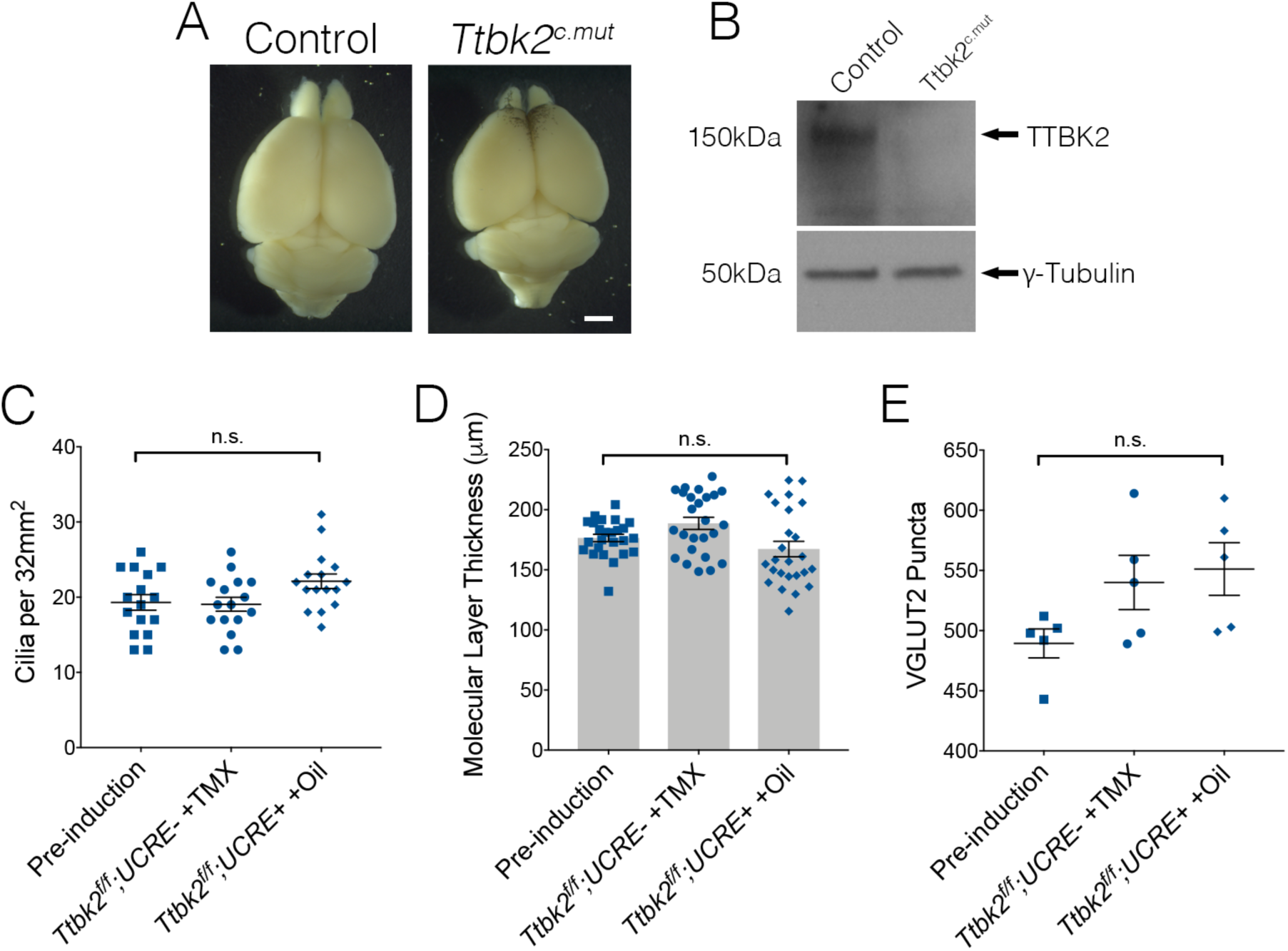
Conditional removal of *Ttbk2* from the adult brain. **(A)** Representative images of Control and *Ttbk2^c.mut^* brains three months after tamoxifen treatment. Scale bar = 1mm **(B)** Western blot analysis of brain lysate from *Ttbk2^c.mut^* animals showing no TTBK2 expressed at three months after tamoxifen injection. **(C-E)** Quantification of cilia abundance **(C)**, molecular layer thickness **(D)**, and VGLUT2 puncta amounts **(E)** across controls. There is no significant difference between control genotypes compared to each other for any of these metrics using a One-way ANOVA with Tukey’s correction (each point represents a field counted, 16 field were counted in total. n=1 animal per genotype, error bars indicate SEM).

**Figure S2.**
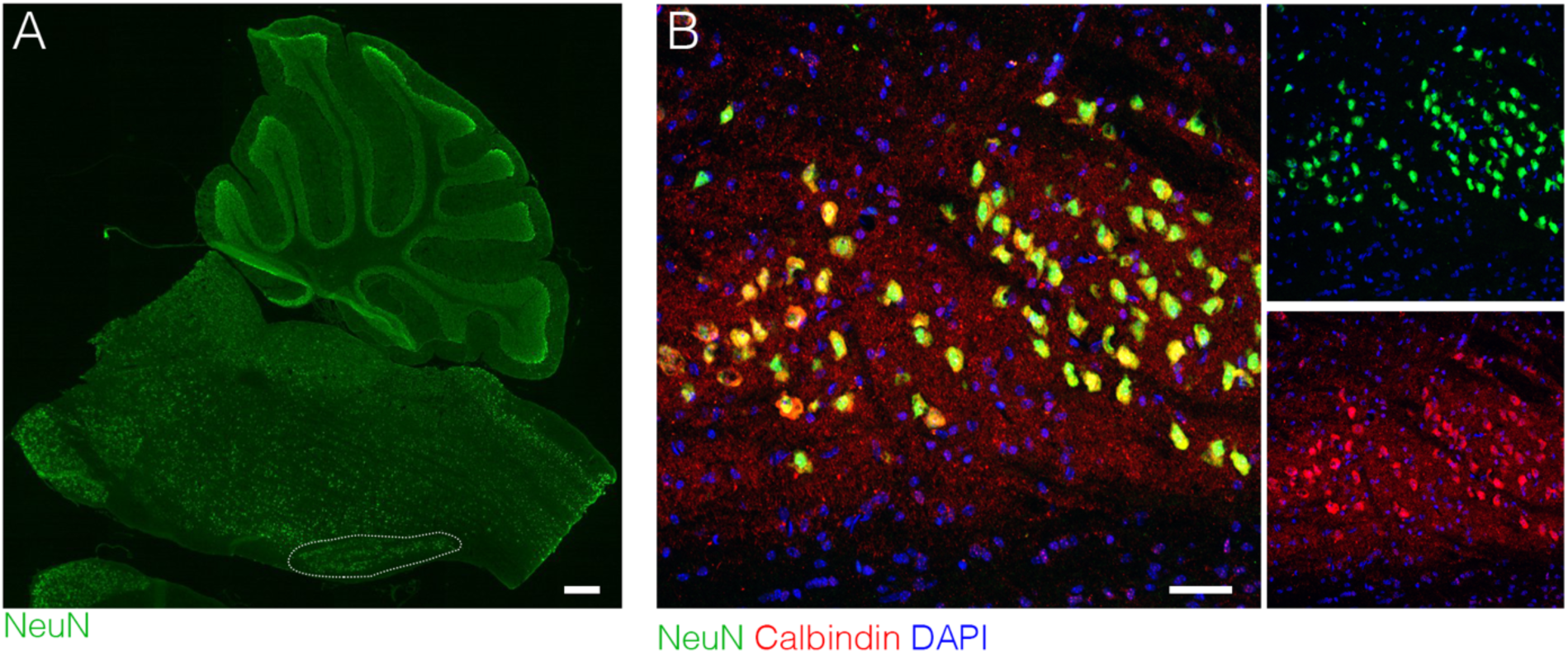
Identification of ION neurons within the medulla. **(A)** The location of the mouse Inferior Olivary Nucleus (ION, dotted outline) within the medulla oblongata of the brain stem. Scale bar = 10mm. **(B)** Neurons within the ION are stained for Calbindin (red), Neun (green), and dapi (blue). The ION is found at the medial ventral part of the medulla oblongata, below the Superior Olivary Nucleus. Scale bar = 100μm.

**Figure S3.**
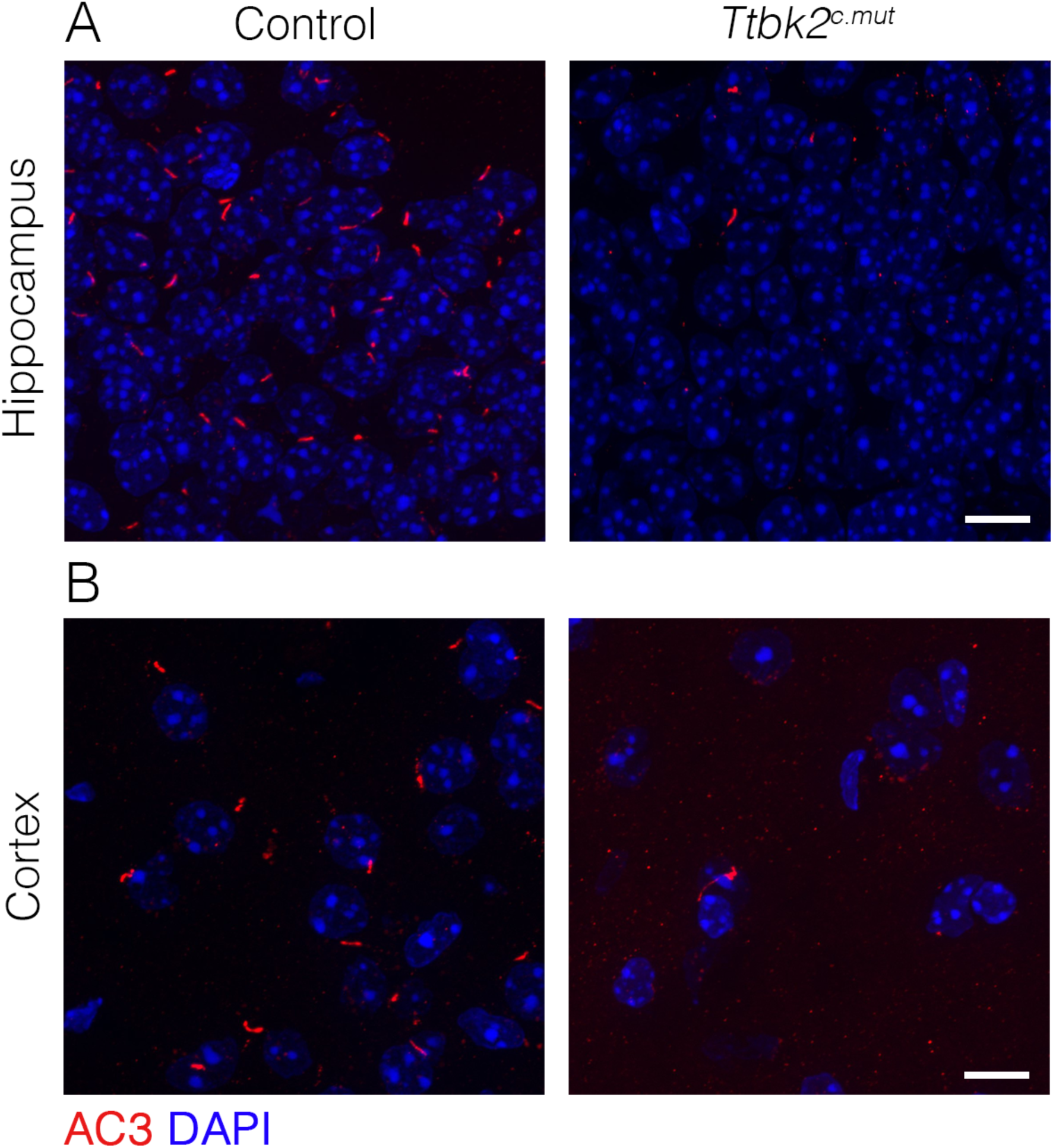
Cilia loss is ubiquitous throughout the brain of *Ttbk2^c.mut^* animals. **(A,B)** Sagittal sections of *Ttbk2^c.mut^* brains stained for AC3 to label cilia (red) and DAPI for nuclei. Neurons in the hippocampus **(A)** and the cortex **(B)** have lost cilia three months after tamoxifen injections in *Ttbk2^c.mut^* animals. Scale bar = 20μm.

**Figure S4.**
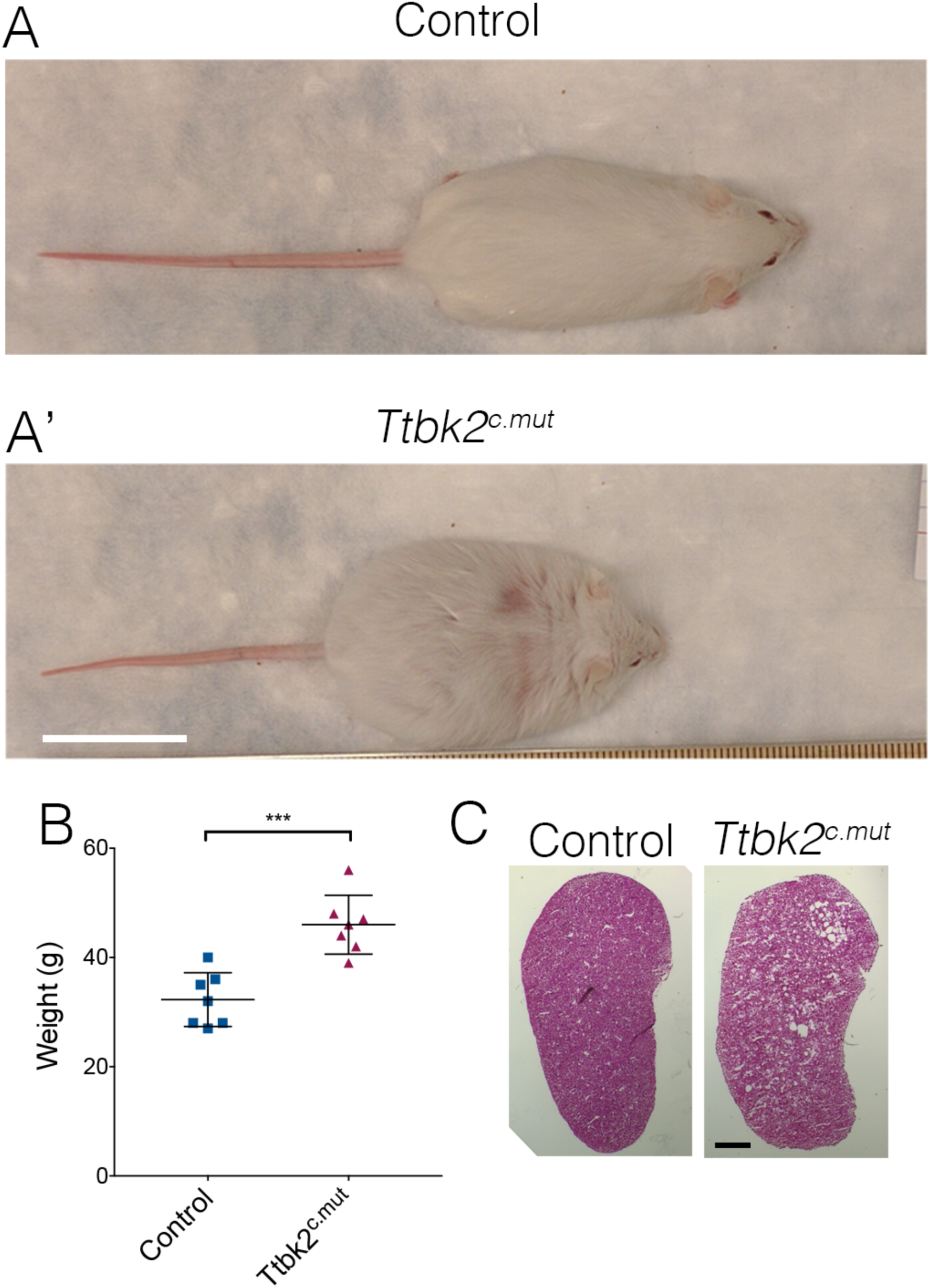
*Ttbk2^c.mut^* animals have phenotypes shared with other ciliopathy models (A-A’) Representative images of Control **(A)** and *Ttbk2^c.mut^* **(A’)** mice three months after tamoxifen treatment. *Ttbk2^c.mut^* mice have gained more weight compared to littermate controls. Scale bar = 1in **(B)** Quantification of weight gain in *Ttbk2^c.mut^* mice compared to Controls. (n=7 biological replicates, p=0.0003). **(C, C’)** H&E staining of kidneys from control **(C)** and *Ttbk2^c.mut^* mice **(C’).** *Ttbk2^c.mut^* mice have polycystic kidneys. Scale bar = 100μm.

**Figure S5.**
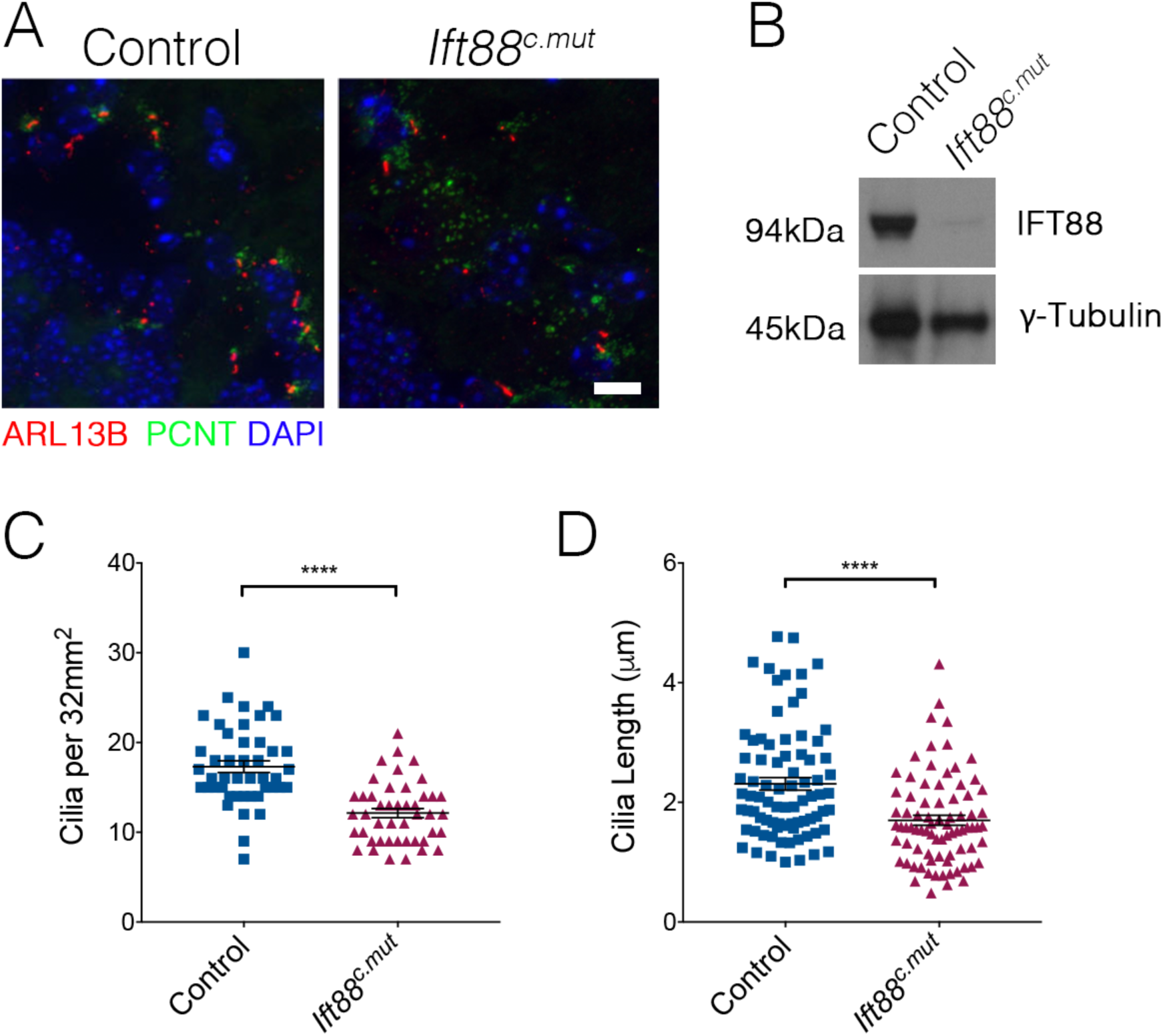
*Ift88^c.mut^* have fewer, shorter cilia throughout the cerebellum. **(A)** Representative images illustrating cilia loss in the cerebellum of *Ift88^c.mut^* animals, immunostained for ARL13B to label cilia (red), PCNT to label centrosomes (green) and nuclei (blue) Scale bar = 20μm. **(B)**Quantification of cilia loss. Unlike in the cerebellum of *Ttbk2^c.mut^* mice, cilia loss is less drastic in the cerebellum of *Ift88^c.mut^* animals (each point represents a field scored, 45 fields scored per genotype. n=3 biological replicates. p<0.0001 by student’s unpaired t-test, error bars indicate SEM 5.18 +/- 0.82) **(C)** Quantification of cilia length between Control and *Ift88^c.mut^.* Cilia in *Ift88^c.mut^* cerebellum are shorter (each point represents a single cilium, 80 cilia were measured between genotypes. n=3 biological replicates, p < 0.0001 by student’s unpaired t-test, error bars indicate SEM -0.6102 +/- 0.1342). **(D)** Western blot analysis of loss of IFT88 in brain lysate of four month old *Ift88^c.mut^* animals.

**Figure S6.**
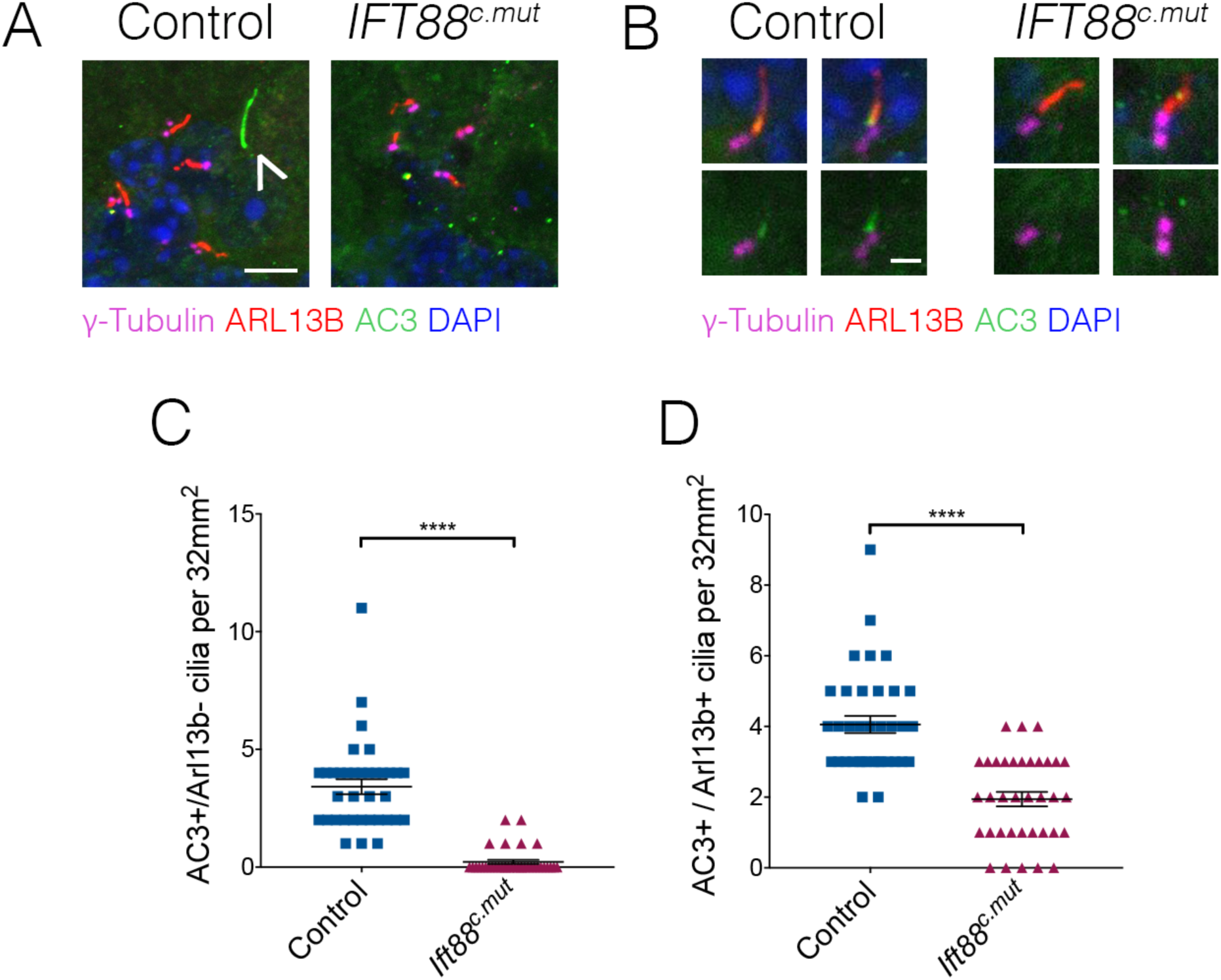
Ciliary composition is perturbed in the brains of *Ift88^c.mut^* animals. (A-B) Cilia from 6 month old Control and *Ift88^c.mut^* stained for γ-Tubulin to label centrosomes (magenta), ARL13B to label cilia membrane (red), AC3 to label cilia membrane (green) and dapi (blue). *Ift88^c.mut^* lose AC3+ cilia (**A**) and a subset of AC3+/ARL13B+ double positive cilia (**B**). Scale bar = 5μm (**A**) and 1μm (**B**). **(C)** Quantification of cilia that are only AC3+ throughout the cerebellum. *Ift88^c.mut^* animals have a strong reduction in this population of cilia. (each point represents a field scored, 36 field scored per genotype. n=3 biological replicates. p<0.0001 by student’s unpaired t-test, error bars indicate SEM -3.19 +/- 0.33) **(D)** Quantification of double positive AC3+/ARL13b+ cilia throughout the cerebellum. Similar to AC3+ data, there is a reduction of this population of cilia in *Ift88^c.mut^* animals (each point represents a field scored, 36 field scored per genotype. n=3 biological replicates. p<0.0001 by student’s unpaired t-test, error bars indicate SEM -2.11 +/- 0.32).

